# Generalization of the minimum covariance determinant algorithm for categorical and mixed data types

**DOI:** 10.1101/333005

**Authors:** Derek Beaton, Kelly M. Sunderland, ADNI, Brian Levine, Jennifer Mandzia, Mario Masellis, Richard H. Swartz, Angela K. Troyer, ONDRI, Malcolm A. Binns, Hervé Abdi, Stephen C. Strother

**Affiliations:** Rotman Research Institute at Baycrest Health Sciences; Baycrest Health Sciences; ADNI; Department of Clinical Neurological Sciences, Western University; University of Toronto; Department of Medicine (Neurology), Sunnybrook Health Sciences Centre; ONDRI; The University of Texas at Dallas; Departments of Psychology and Medicine (Neurology), University of Toronto

**Keywords:** Minimum covariance determinant, correspondence analysis, categorical data, outliers, robust, neuroinformatics

## Abstract

The minimum covariance determinant (MCD) algorithm is one of the most common techniques to detect anomalous or outlying observations. The MCD algorithm depends on two features of multivariate data: the determinant of a matrix (i.e., geometric mean of the eigenvalues) and Mahalanobis distances (MD). While the MCD algorithm is commonly used, and has many extensions, the MCD is limited to analyses of quantitative data and more specifically data assumed to be continuous. One reason why the MCD does not extend to other data types such as categorical or ordinal data is because there is not a well-defined MD for data types other than continuous data. To address the lack of MCD-like techniques for categorical or mixed data we present a generalization of the MCD. To do so, we rely on a multivariate technique called correspondence analysis (CA). Through CA we can define MD via singular vectors and also compute the determinant from CA’s eigenvalues. Here we define and illustrate a generalized MCD on categorical data and then show how our generalized MCD extends beyond categorical data to accommodate mixed data types (e.g., categorical, ordinal, and continuous). We illustrate this generalized MCD on data from two large scale projects: the Ontario Neurodegenerative Disease Research Initiative (ONDRI) and the Alzheimer’s Disease Neuroimaging Initiative (ADNI), with genetics (categorical), clinical instruments and surveys (categorical or ordinal), and neuroimaging (continuous) data. We also make R code and toy data available in order to illustrate our generalized MCD.

## Introduction

Today’s large scale studies contain a wide array of data from multiple sources that span a variety of data types. For examples, certain types of genetic data (single nucleotide polymorphisms) are categorical, surveys and clinical instruments are often ordinal, and other measures, such as reaction times or neuroimaging, are continuous. Such data sets are typically open or broadly accessible, and thus analyzed by hundreds or thousands of researchers. With many analyses at stake, it is critical to ensure the integrity and accuracy of data as well as the robustness of results. In the Ontario Neurodegenerative Disease Research Initiative (ONDRI), we employ a quality control step as part of the release process to identify anomalies (Sunderland et al., 2019). Our quality control step uses a common and reliable outlier detection technique called the minimum covariance determinant (MCD; Hubert & Debruyne, 2010), which sees use across numerous disciplines (Hadi, Imon, & Werner, 2009; Leys, Klein, Dominicy, & Ley, 2018; Magnotti & Billor, 2014; Verity et al., 2017). However, the MCD can only identify outliers in continuous data. In ONDRI—and many studies like ours—we have data sets comprised entirely of categorical or ordinal data, and complex data sets with mixtures of data types. Clearly we need an outlier detection technique that can accommodate various data types and, preferably, work in much the same way the MCD technique works.

The MCD produces robust estimates for multivariate location (mean) and scatter (covariance). Given a matrix of observations (rows) and variables (columns), the MCD finds a subset of observations that have minimum scatter. In the MCD, scatter is defined by the determinant, and thus the search for a minimum determinant from a (covariance) matrix. Because the MCD algorithm searches for the smallest scatter of observations, it also helps identify multivariate outliers via robust Mahalanobis distances. Since the introduction of the Fast-MCD approach (Rousseeuw & Van Driessen, 1999), there have been many improvements and variations on the technique, such as robust PCA (Hubert, Rousseeuw, & Branden, 2005), deterministic MCD (Hubert, Rousseeuw, & Verdonck, 2012), and extensions to address high-dimensional data through regularization (Boudt, Rousseeuw, Vanduffel, & Verdonck, 2017). See Hubert, Debruyne, and Rousseeuw (2017) for an overview on the MCD and various extensions thereof. Though the MCD and its variants are standard techniques to identify both robust covariance structures and outliers, MCD approaches work only for data assumed to be continuous. So, how can we adapt MCD approaches to non-quantitative data? The major barrier to adapt the MCD for non-quantitative data is because there are no standard definitions of Mahalanobis distances for non-quantitative data.

Goodall (1966) noted that there are generally two issues with the application of standard computations and rules of Mahalanobis distances to non-quantitative data: (1) “If […] quantitative and binary attributes are included in the same index, these procedures will generally give the latter excessive weight.” (p. 883), and (2) “indices appropriate to quantitative [data have] been applied to ordered, non-quantitative attributes […] by arbitrarily assigning metric values to their different ordered states.” (p. 883). Clearly, we do not want categorical data to provide undue influence on Mahalanobis distance estimates, nor should we arbitrarily apply metric values to categorical or ordinal data. While there are many Mahalanobis-like distances or alternative measures of similarity for non-quantitative data (Bar-hen & Daudin, 1995; Bedrick, Lapidus, & Powell, 2000; Boriah, Chandola, & Kumar, 2008; Leon & Carrière, 2005; McCane & Albert, 2008) many still have the drawbacks noted by Goodall (1966). If we want to develop a MCD algorithm for non-quantitative data, we need to define a Mahalanobis distance that neither imposes undue influence nor arbitrarily assigns values.

Here we introduce an approach that generalizes the MCD for use with a variety of data types: categorical, ordinal, continuous, and mixtures thereof. Our generalization of the MCD is possible through the use of Correspondence Analysis (CA). By way of CA, we can obtain both the determinant of a matrix from its eigenvalues, and Mahalanobis distances from its singular vectors. Our paper is outlined as follows. We first show how the SVD provides the two key requirements for the MCD algorithm: the determinant of a covariance matrix and Mahlanobis distances. Then, we show how we can generalize these properties of the SVD in order to compute Mahalanobis distances for categorical data through CA. With Mahalanobis distances for categorical data, we can then formalize a generalized MCD for categorical data via Multiple CA. Specifically, our generalized MCD makes use of a technique called “subset multiple correspondence analysis”. Next we illustrate our method on a toy (simulated) genetics dataset of categorical variables. We then show how our generalized MCD works at scale. First we use the MCD for survey data from ONDRI. Because survey data can be treated as either categorical or ordinal data, we apply our generalized MCD to the same survey data under categorical and ordinal assumptions. Coding the survey data both ways allows us to compare and contrast the behavior of our generalized MCD under each condition. We then apply our generalized MCD to a mixed data set from the Alzheimer’s Disease Neuroimaging Initiative (ADNI). In the ADNI example we show how our generalized MCD extends to mixed data with categorical (genetics), ordinal (a clinical instrument), and continuous (brain volumes) data. In the ADNI example we highlight particular outliers identified from our generalized MCD, and then show how to understand why those individuals are outliers. Finally we discuss the technique with respect to the results, as well as provide future directions, limitations, and concluding remarks for our generalized MCD.

## A Generalization of the MCD for non-quantitative data

### Software and notation

We used the `papaja` package (Aust & Barth, 2018) to write this manuscript via RMarkdown. We make our software, code, and some examples of the GMCD available via the *Ou*tliers and *R*obust *S*tructures (OuRS) package at https://github.com/derekbeaton/ours.

Bold uppercase letters denote matrices (e.g., **X**), bold lowercase letters denote vectors (e.g., **x**), and italic lowercase letters denote specific elements (e.g., *x*). Upper case italic letters denote cardinality, size, or length (e.g., *I*) where a lower case italic denotes a specific index (e.g., *i*). A generic element of **X** would be denoted as *x*_*i,j*_. Common letters of varying type faces for example **X, x**, *x*_*i,j*_ come from the same data struture. Vectors are assumed to be column vectors unless otherwise specified. Two matrices side-by-side denotes standard matrix multiplication (e.g., **XY**), where ⊙ denotes element-wise multiplication. The matrix **I** denotes the identity matrix. Superscript ^*T*^ denotes the transpose operation, superscript ^−1^ denotes standard matrix inversion, and superscript ^+^ denotes the Moore-Penrose pseudo-inverse. Recall that for the generalized inverse: **XX**^+^**X** = **X**, (**X**^+^)^+^ = **X**, and (**X**^*T*^)^+^ = (**X**^+^)^*T*^. The diagonal operation, diag{}, when given a vector will transform it into a diagonal matrix, or when given a matrix, will extract the diagonal elements as a vector. Say **D** is some diagonal matrix and recall that the transpose of a diagonal matrix is itself and thus **D**^*T*^ = **D**. We denote ⌊.⌋ as the floor function. We use det{} to indicate the determinant of a square symmetric matrix. Finally, we reserve some letters to have very specific meanings: (1) **F** denotes component (sometimes called factor) scores where, for example, **F**_*I*_ denotes the component scores associated with the *I* items, (2) **O** and **E** mean “observed” and “expected”, respectively, as used in standard Pearson’s *X*^2^ analyses, where for example **O**_**X**_ denotes the “observed matrix derived from **X**”, (3) **Z** means a centered and/or normalized matrix derived from another, e.g., **Z**_**X**_ is a centered and/or normalized version of **X**, and (4) blackboard bold letters mean some *a priori* or known values, for example 𝔼**X** would reflect some previously known expected values associated with **X** but not necessarily derived from **X**. From this point forward we also refer to Mahalanobis distance as MD.

### Overview of MCD algorithm

Given a matrix **X** with *I* rows and *J* columns where *J* < *I* and **X** is full rank. The MCD requires an initialization of subset of size *H*—a random subsample from *I* rows of **X**—where ⌊(*I* + *J* + 1)*/*2⌋ ≤ *H* ≤ *I*. From **X**_*H*_ we compute the column means as ***µ***_*H*_. The column-wise centered (on ***µ***_*H*_) version of **X**_*H*_ is denoted as **Z**_*H*_. The MCD algorithm works as follows:

1. Compute the covariance matrix 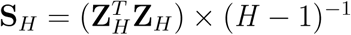.
2. Compute the determinant of **S**_*H*_.
3. Compute the squared MD for each observation in **X** as 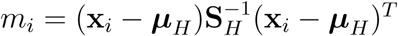.
4. Set **X**_*H*_ as the subset of *H* observations with the smallest MDs in Step 3.

The size of *H* is controlled by *α* to determine the subsample size; *α* exists between 0.5 and 1. The above steps are repeated until we find a minimum determinant either through optimal search or within a specified number of iterations. We denote the *H* subset with the minimum determinant as Ω. For the Ω subset, we obtain a robust mean vector ***µ***_Ω_, a robust covariance 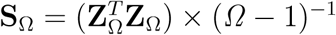, and robust square MDs for each *i* observation from **X** as 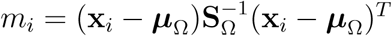.

The two key elements for MCD algorithms are: (1) the determinant of the subsample covariance matrix, which helps identify the smallest scatter of observations and (2) the squared MDs for all observations derived from the subsample covariance matrix, which both helps identify outliers and aids in the search for a minimum determinant. Both the determinant of a matrix and the squared MDs for observations can be computed from the singular value decomposition (SVD): (1) the determinant can be computed as the geometric mean of the eigenvalues (Boudt et al., 2017; SenGupta, 1987) and (2) squared MD can be computed as the sum of the squared row elements of the singular vectors (Barkmeijer, Bouttier, & Van Gijzen, 1998; Brereton, 2015; Stephenson, 1997). In the following sections we first show how to compute what is needed (i.e., determinants, squared MDs for subsamples and out-of-sample data) only through the SVD, which provides the basis of how to generalize the MCD to additional data types.

### Mahalanobis distances via PCA for continuous data

Given **X** with *I* rows and *J* columns, and have **Z**_**X**_ as the centered and/or normalized version of **X**. Assume that **Z**_**X**_ is full rank, where *L* is the rank of **Z**_**X**_ and *L* ≤ *J* < *I*. First we define the sample covariance matrix of **X** as 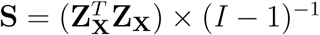. Squared MD is **m** = (**Z**_**X**_**S**^−1^ ⊙ **Z**_**X**_)**1** where **1** is a conformable *J* × 1 vector of 1s and **m** is a *I* × 1 vector of squared MDs. If we relax the definition and exclude the degrees of freedom scaling factor (*I* − 1):

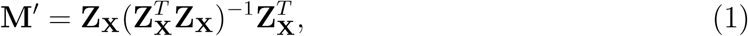

where **m**′ = diag{**M**′} and **m**′ = **m** × (*I* − 1)^−1^. Henceforth we refer to squared MD as 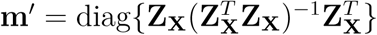. Note that 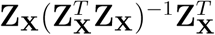 is also called the “orthogonal projection matrix” (Yanai, Takeuchi, & Takane, 2011), and that 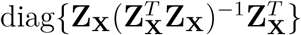 is equal to the influence measure of “leverage” in regression and PCA (Mejia, Nebel, Eloyan, Caffo, & Lindquist, 2017; Wold, Esbensen, & Geladi, 1987). We can also define squared MDs by way of principal components analysis (PCA). We can perform the PCA of **X** through the SVD as:

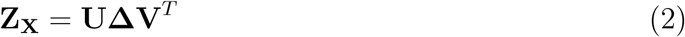

where (1) **U** and **V** are orthonormal left and right singular vectors of sizes *I* × *L* and *J* × *L*, respectively with **U**^*T*^ **U** = **I** = **V**^*T*^ **V** and **U**^*T*^ = **U**^+^ and **V**^*T*^ = **V**^+^, and (2) **Δ** is the *L* × *L* diagonal matrix of singular values and **Λ** = **Δ**^2^ which is a diagonal matrix of eigenvalues (squared singular values). With Eq. (2) we can compute the determinant from the eigenvalues, and then the squared MDs as:

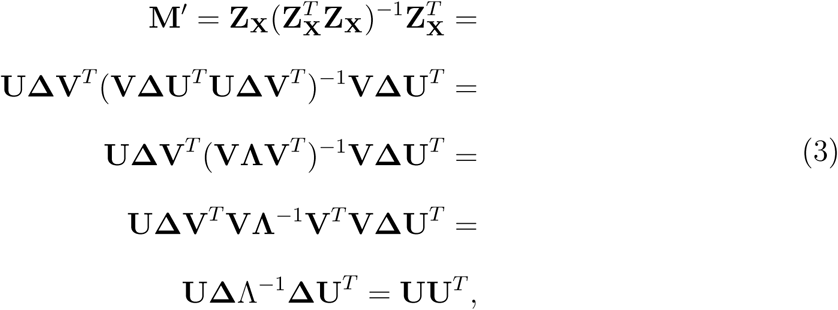

where 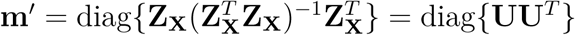. In PCA, component scores are scaled (by the singular values) versions of the vectors with **F**_*I*_ = **UΔ** defined as row component scores. We can also define row component scores through projection (or rotation) as:

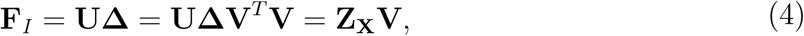

where 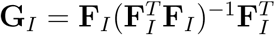 and each row item’s leverage is an element in **g**_*I*_ = diag{**G**_*I*_}. If we expand and substitute **UΔ** for **F**_*I*_, we see that **G**_*I*_ = **UΔ**(**ΔU**^*T*^ **UΔ**)^−1^**ΔU**^*T*^ = **UU**^*T*^, thus **m**′ = **g**_*I*_ = diag{**UU**^*T*^}.

MCD algorithms find a subsample of observations that has a minimum determinant and produces a robust covariance structure. From this robust structure, squared MDs are computed for both the subsample and the excluded—or out of sample—observations. We can compute the squared MDs on the subsample and out of sample observations through projection of data onto the space spanned by a given set of components. First, we refer to a *K* × *J* matrix **Y** as a new set of observations with the same variables (columns), and preprocessed in the same way as **X**. Thus we have **Z**_**Y**_ which has been, for example, centered by the column-wise center of **X**. For **Z**_**Y**_, the squared MDs are 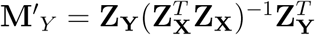. Alternatively, we can project **Z**_**Y**_ onto previously defined components, as established in Eq. (4) To compute component scores for **Z**_**Y**_ with respect to the PCA of **Z**_**X**_ as **F**_*K*_ = **Z**_**Y**_**V**. Through that projection, we can compute the squared MDs for **Z**_**Y**_ with only **V** and **Δ** from the SVD of **Z**_**X**_ as defined in Eq. (2) as

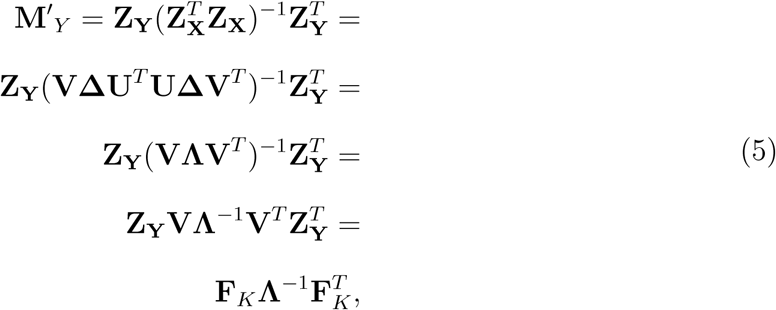

where the squared MDs are 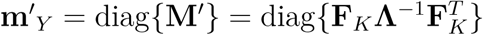. Now let us assume that **Y** is comprised of both included and excluded observations, or in terms of subsamples: the *H* included samples and the 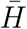 excluded samples. Then 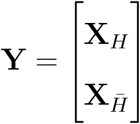. In this case, **Z**_**Y**_ is, for example, centered on the column-wise mean of **X**_*H*_. We can compute squared MDs for both the *H* included and the 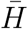 excluded samples as 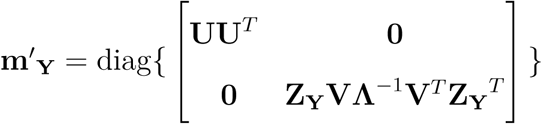.

Next we show how to use the SVD under different assumptions and metrics so that we can define a MCD algorithm for various data types. We show how to do so through an analog of PCA designed for categorical data called multiple correspondence analysis.

### Multiple correspondence analysis

Multiple correspondence analysis (MCA) generalizes both (1) PCA to categorical data (with observations on the rows) and (2) standard correspondence analysis (CA) from two-way contingency to N-way contingency tables (Abdi & Valentin, 2007; Greenacre, 1984; Greenacre & Blasius, 2006; Lebart, Morineau, & Warwick, 1984). However, MCA can accommodate many other data types – which we cover in detail in the next section. MCA operates under the assumption of independence (i.e., *χ*^2^).

Say we have an *I* × *A* matrix called **A**, as in Table 1a, where the rows are observations and the columns are categorical variables where each cell contains a nominal level of each variable. These data can be represented in complete disjunctive coding—as in Table 1b—where each variable is represented by a vector of length *N*_*a*_, where *N*_*a*_ is the number of levels or nominal values for the *a*^th^ variable. We refer to the disjunctive of **A** as an *I* × *J* matrix called **X**.

**Table 1.** Example categorical and disjunctive data.

|  | Var. 1 | Var. 2 |
| --- | --- | --- |
| *Subj.1* | B | YES |
| *Subj.2* | A | YES |
| ... | ... | ... |
| *Subj.I-1* | A | NO |
| *Subj.I* | C | YES |

(b) *Disjunctive representation of two variables.*
|  | Var. 1 |  |  | Var. 2 |  |
| --- | --- | --- | --- | --- | --- |
|  | A | B | C | YES | NO |
| *Subj.1* | 0 | 1 | 0 | 1 | 0 |
| *Subj.2* | 1 | 0 | 0 | 1 | 0 |
| ... | ... | ... | ... | ... | ... |
| *Subj.I-1* | 1 | 0 | 0 | 0 | 1 |
| *Subj.I* | 0 | 0 | 1 | 1 | 0 |

*Note.* Hypothetical example of categorical data. (a) shows two variables Variable 1 and Variable 2 with 3 and 2 levels respectively. (b) shows the disjunctive code for these data. Note that the sum of each row is equal to the total number of original variables, and that the sum within each set of disjunctive columns per variable is equal to the total number of rows. The column sums indicate the frequency of a particular level within the data.

MCA requires specific preprocessing and constraints applied to the rows and columns in order to decompose a matrix under the assumption of independence. To perform MCA, we first compute the row and column weights:

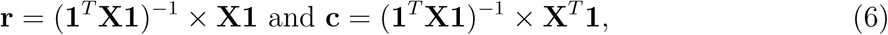

where **r** and **c** are the row and column marginal sums, respectively, divided by the total sum. In the case of purely disjunctive data, each value in **r** is identical as each row contains the same number of 1s (see Table 1b): Each element in **r** is simply the total number of columns in **A** divided by the sum of all elements of **X**. Similarly, **c** contains the column sums of **X** divided by the total sum. Next we compute observed (**O**_**X**_) and expected (**E**_**X**_) matrices of **X** as **O**_**X**_ = (**1**^*T*^ **X1**)^−1^ × **X** and **E**_**X**_ = **rc**^*T*^, respectively, and deviations computed as **Z**_**X**_ = **O**_**X**_ − **E**_**X**_. MCA is performed with the generalized SVD (GSVD) of **Z**_**X**_:

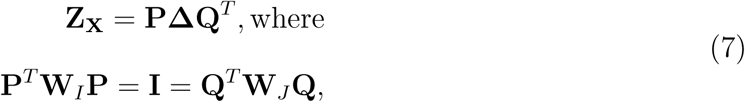

and where **W**_*I*_ = diag{**r**}^−1^ and **W**_*J*_ = diag{**c**}^−1^. The *generalized* singular vectors are computed as 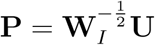 and 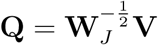. We obtain the results of the GSVD of **Z**_**X**_ through the SVD of 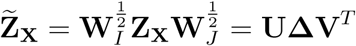 as in Eq. (2). We can see the relationship between **Z**_**X**_ and 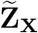 via substitution:

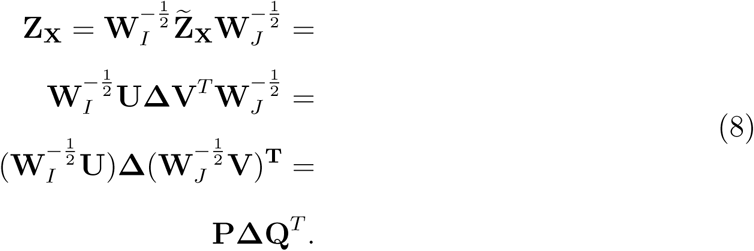

For simplicity, we refer to the GSVD where needed in triplet notation as GSVD(**W**_*I*_, **Z**_**X**_, **W**_*J*_) with row constraints (e.g., **W**_*I*_), data (e.g., **Z**_**X**_), and column constraints (e.g., **W**_*J*_). To note, we have taken liberty with the standard GSVD triplet notation (see Holmes, 2008) and present the triplet more akin to its multiplication steps.

#### A computational note on MCA

Because of the way we represent data for MCA—see the disjunctive coding example in Table 1—and because of how the SVD works, MCA produces “null dimensions” (Greenacre, 1984). These “null dimensions” are produced through the use of the SVD (or eigendecomposition) where the singular (or eigen) vectors are undetermined and the singular (or eigen) values are effectively zero. These “null dimensions” are to be discarded. In all subsequent sections, our use of the MCA, CA, and related methods also excludes these “null dimensions”. In effect, we only retain the dimensions with non-zero eigenvalues.

### Mahalanobis distances via MCA for categorical data

We showed for continuous data that we can obtain the squared MD from the diagonal of the crossproduct of left singular vectors as diag{**UU**^*T*^}. We use the same premise to define a squared MD for categorical data by way of MCA. MCA is performed as GSVD(**W**_*I*_, **Z**_**X**_, **W**_*J*_) where we apply the SVD as 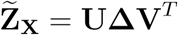 (see Eqs. (7) and (8). Recall that 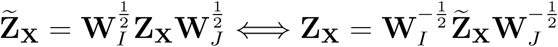 where **Z**_**X**_ = **PΔQ**^*T*^ (see Eq. (8)). In the case of 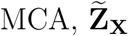 would be the analog of **Z**_**X**_ in PCA (see Eq. (2)), thus via the SVD we still have 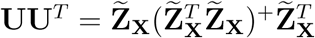. We replace the standard inversion with the pseudo-inverse—and only use non-null dimensions—because of the inherent collinearity in 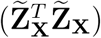. Furthermore 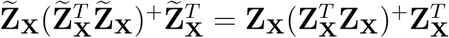, which leads to

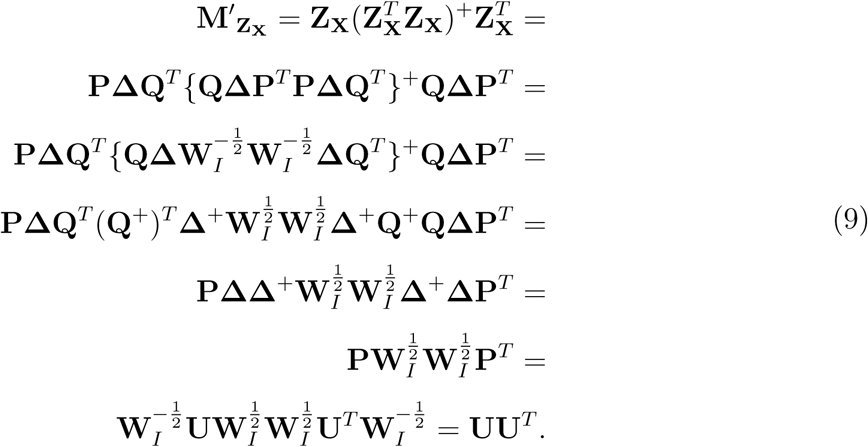

Thus 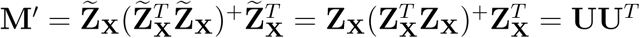 is a squared MD for categorical data via MCA, à la Eqs. (1) and (3). Because of the way we define squared MD here—the singular vectors for the observations (i.e., **U**)—there exists no ambiguity nor arbitrary decisions in the definition of squared MD for categorical data (à la, Goodall, 1966).

We can compute MCA row component scores as **F**_*I*_ = **W**_*I*_**PΔ** or through projections. For MCA component scores through projections, it is easier to use “profile” matrices where **Φ**_*I*_ = **W**_*I*_**O**_**X**_ are the row profiles. For profile matrices, each element in **X** divided by its respective row sum for the row profile matrices. MCA row component scores via projection are:

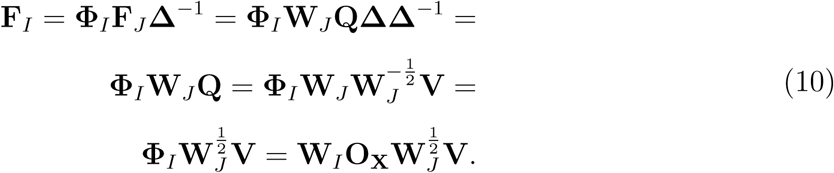

Note that 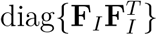 are referred to as *χ*^2^-distances. We can compute sqaured MDs through row component scores where 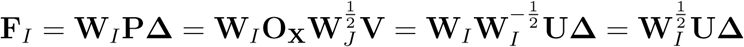 and so:

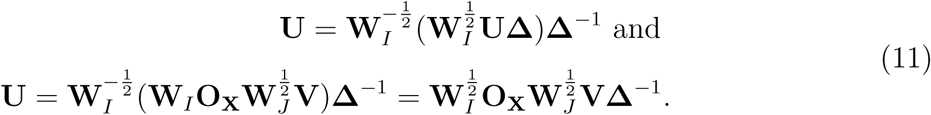

While Eq. (10) and Eq. (11) suggest ways to compute squared MD for sub- or out-of sample categorical data—akin to Eq. (5)—there exists a major barrier to the computation of *correct* squared MD for sub- or out-of sample categorical data. The sub-sampling procedure cannot be performed on the original categorical data **A** nor on the initial disjunctive table **X**. We cannot subsample **A** and **X** because levels within categorical variables will likely be excluded in the subsample. Exclusion of levels produces incorrect (i.e., lower) estimates of squared MDs for the excluded subsample. To correctly compute MDs, we need to maintain the representation of all levels. To do so, we require a specific form of MCA called subset MCA (Greenacre, 2017; Greenacre & Pardo, 2006), which is a specific case of Generalized CA (Escofier, 1983, 1984).

### Escofier’s Generalized Correspondence Analysis and Subset MCA for the MCD

Hiding in the broader CA literature is “generalized correspondence analysis” (GCA: Escofier, 1983, 1984)—as referred to by Grassi and Visentin (1994). Escofier established the idea of GCA as “(DATA – MODEL) / MARGINS”. GCA has many uses well beyond the standard applications in CA and MCA such as for missing data and assumptions that deviate from the typical assumptions of independence (*χ*^2^) for CA (e.g., quasi-CA with “quasi-independence” and “quasi-margins”; Leeuw & Heijden, 1988; Van der Heijden, De Falguerolles, & Leeuw, 1989). With a complete disjunctive data set **X**, for the GCA perspective of MCA, we characterize each cell of 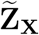 with the generic element 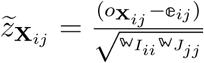. The model (𝔼) and the margins (𝕎_*I*_ and 𝕎_*J*_), could be defined in almost any way; it is not required to compute the model (expected values) nor margins from the data. To help simplify the notation, we can use the GSVD to represent the GCA approach: ℤ_**X**_ = **O**_**X**_ − 𝔼 and GSVD(𝕎_*I*_, Z_**X**_, 𝕎_*J*_) (see Eqs. (7) and (8)). For our problem, we only need a simple change to the generic element 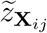. First we compute all matrices required for MCA exactly as initially defined in Eqs. (7) and (8): **O**_**X**_, **E**_**X**_, **r** and **c**.

Let us assume that **O**_**X**_, **E**_**X**_, and **r** are constructed as partitions defined by the *H* and 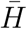 subsets as: 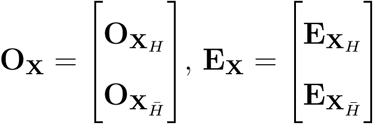, and 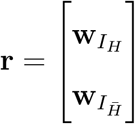. To maintain the structure of the data and to be able to perform the necessary steps for a generalized MCD we define a generalized expected matrix as 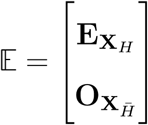; that is, we replace part of the expected matrix with part of our observed matrix. Our assumption is that the expected values are the observed, thus 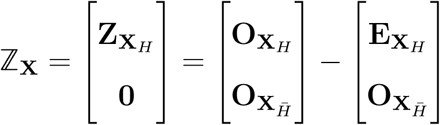. Any subsample of *H* and 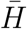 guarantee that we maintain the same shape (rows and columns) as **X** when we use 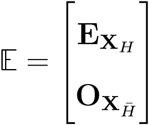.

Like in Eqs. (7) and (8), we have 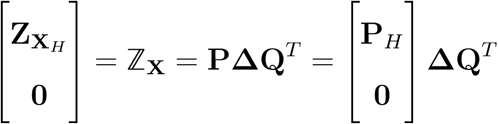, where **P**^*T*^ **W**_*I*_**P** = **I** = **Q**^*T*^ **W**_*J*_**Q** and 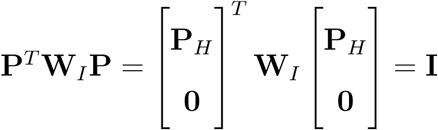. Furthermore, recall that 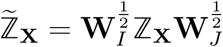 and thus 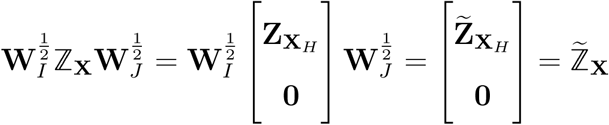. Just as in Eq. (8) we would apply the SVD as:

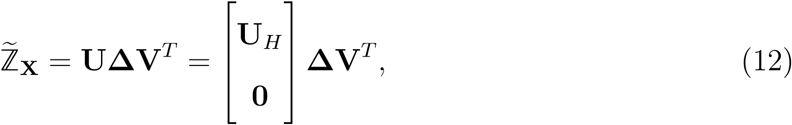

where **U**^*T*^ **U** = **I** = **V**^*T*^ **V** and 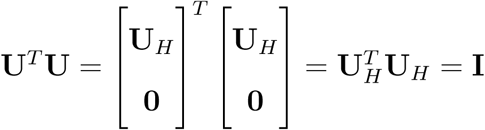.

We can simplify this approach through a specific application of GCA called “subset MCA” (Greenacre, 2017; Greenacre & Pardo, 2006; Hendry, North, Zewotir, & Naidoo, 2014). To compute subset MCA for use in the MCD we use the matrices defined in Eqs. (7) and (8). Specifically 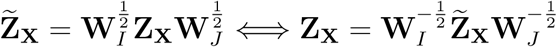. In subset MCA we require 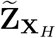 which is the *H* subsample of 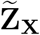 and perform the SVD as

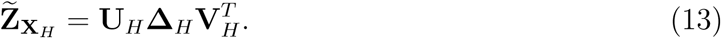

Which leaves us with the complement 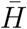 where 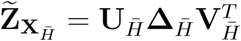.

We can compute the determinant for *H* subsamples via the geometric mean of the eigenvalues 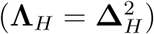 and we can compute robust squared MDs in subset MCA with the row profiles as in Eqs. (10) and (11). We compute the component scores for the full *I* set from subset MCA as

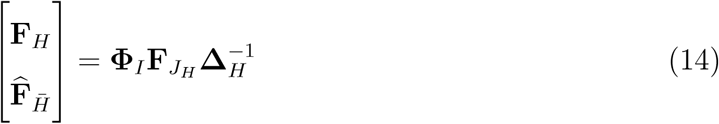

which allows us to compute 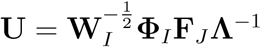 because 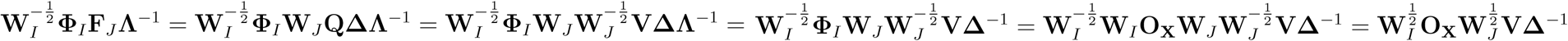. We can compute the squared MDs for the analyzed and excluded subsamples from the subset MCA results as:

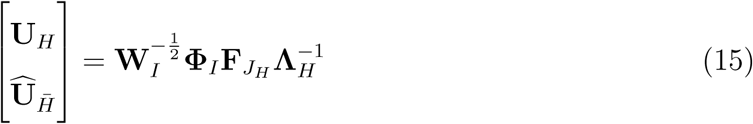

where the **U**_*H*_ in Eq. (15) is exactly the **U**_*H*_ in Eqs. (12) and (13). Alternatively we can compute Eq. (15) from (14) as 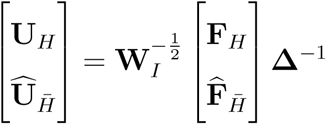, and thus squared MDs are computed as 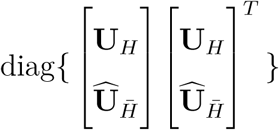.

### MCD algorithm for categorical data

Here we outline the core steps required for a MCD via subset MCA. First the MCD requires an initial random subsample of size *H* where ⌊(*I* + *J* + 1)*/*2⌋ ≤ *H* ≤ *I*. The *H* subset of 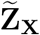 is denoted as 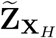.

1. Apply the SVD: 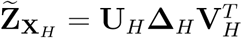
2. Compute the determinant of 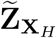 as the geometric mean of **Λ**_*H*_
3. Compute the squared MDs for the full *I* sample as 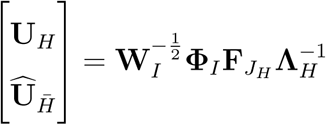
4. Set 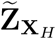 as the subset of *H* observations with the smallest squared MDs computed from Step 3.

These steps are repeated until a we find a minimum determinant (Step 2) or stop at a given number of iterations. Like in the traditional MCD, Step 3 helps find a minimum determinant because the subset of *H* observations with the smallest squared MDs produce a smaller or equal determinant in subsequent iterations.

#### Differences between traditional MCD and our proposed MCD

Primarily, our approach to the generalization of a MCD for categorical data maintains the two most important features of the MCD algorithm: (1) search for a minimum determinant to identify a robust covariance structure and (2) compute robust squared MDs with respect to that structure, in order to identify outliers. In effect, like with the standard MCD approach, ours also helps in diagnostic steps that could be susceptible to masking effects. However there are some key differences between our approach to a MCD algorithm for categorical data and the traditional MCD (which is, generally, only defined for continous data). These differences stem from our approach via MCA and the constraints imposed on the problem by categorical data. The most considerable issue with categorical data is that rare response levels are likely sources of an individual’s outlierness. But those levels cannot be removed, else the individuals with those response levels end up with *smaller* MDs simply because those rare levels are no longer represented. We retain all the key computational aspects, while relaxing some of the theoretical.

First, our approach uses what should be considered a pseudo-determinant. Both 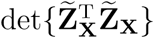 and 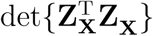 are by definition zero because the multiple columns used to represent a single variable within the disjunctive table (see Table 1) are linear combinations of one another. Computationally, we use the full set of non-null dimensions to compute the pseudo-determinant (i.e., all components with non-zero eigenvalues). Related to this is that it is possible for different subsets, in the search for a minimum determinant, to have different dimensionality; specifically, subsets could span fewer dimensions than the original data. However in practice, this is not much of an issue because fewer dimenions does not necessarily produce a smaller pseudo-determinant (i.e., geometric mean of the eigenvalues for the existing dimensions). With respect to dimensionality in MCA, the “true” dimensionality is typically much smaller than just the non-null dimensions (i.e., effectively zero eigenvalue). Typically in MCA, a correction is applied to retain fewer dimensions and then adjust the eigenvalues through, for example, the Benzecri (Benzécri, 1979) or Greenacre (Greenacre, 2017) corrections. While this is typical for MCA, we recommend against this for our MCD procedure. Generally, the “masking effect” occurs because of anomalous values in the smaller dimensions (i.e., typically dimensions with small eigenvalues).

In the traditional MCD, a new center is computed for each *H* subsample in order to center that *H* subsample on its respective column means. In turn, the subsample identified for the minimum determinant produces a “robust center”. Our generalized approach to MCD does not produce a robust center, per se. Rather, via the search procedure in our approach, we detect both the center and minimum scatter (smallest determinant) from a set of points *within the existing space*, as opposed to defining a new space from the subsample. As previously noted, this is a limitation imposed by categorical data: if we were to drop levels of variables, then the MD of individuals with those levels goes down; this is because those individuals would have exclusively zeros in the disjunctive table. However there is one way—under certain conditions—where a robust center could be computed. A robust center—i.e., the expected matrix **E**_**X**_—could be computed from different *row weights*, say, from only the rows included in the *H* subsample. However, this is only possible (or relevant) when the rows (observations) have different weights. In our approach, all rows have weights of 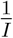. Even with different row weights, our approach leverages the subset CA procedure and thus still searches for a center and minimum scatter within an existing space (defined under assumptions of independence).

## Examples, applications, and extensions

We provide three sets of examples to illustrate the use of our generalized MCD. Because our approach is based on CA, we can capitalize on particular properties of CA to allow for the analyses of non-categorical data such as ordinal, continuous, or mixed (“heterogeneous”) data. We show how to accomodate different data types—including mixed—through a data recoding technique called data “doubling” (a.k.a. fuzzy coding, bipolar coding, or “Escofier transform”). We first illustrate our generalized MCD with a small toy genetic data set of single nucleotide polymorphisms (SNPs; see supplemental section from Beaton, Dunlop, and Abdi (2016)) which are entirely categorical. Next we analyze the miniature survey of autobiographical memory (MSAM; adapted from Palombo, Williams, Abdi, & Levine, 2013), from the Ontario Neurodegernative Disease Research Initiative (ONDRI). While the MSAM is comprised entirely of ordinal responses, we apply our approach to two versions of the MSAM: (1) where each question is treated as a categorical variable, and (2) an approach that preserves the ordinality of the data. We compare and contrast the results from these applications to highlight how GMCD results change. Finally, we provide an analysis of a data set from the Alzheimer’s Disease Neuroimaging Initiative (ADNI) that spans numerous types: categorical data (SNPs), ordinal data (clinical dementia rating), and continuous data (volumetric estimates from brain regions).

### Toy data

The toy data are comprised of SNPs, which are categorical variables. The toy data available as part of the `OuRS` package: https://github.com/derekbeaton/ours. Each SNP contains three possible genotypes: “AA”, “Aa”, and “aa” which are the major homozygote, the heterozygote, and the minor homozygote, respectively. In this example, we set *α* = 0.5.

The toy data are of size *I* = 60 × *C* = 9 (observations by variables). These data were transformed into complete disjunctive coding (as in Table 1) that are of size *I* = 60 × *J* = 25 (observations by columns); note that some SNPs have 3 observed genotypes where as others only have 2 observed genotypes. Figure 1 shows GMCD applied to the toy data set over the course of 1, 000 iterations. Figure 1a shows the search for a minimum determinant (a decrease over iterations). In Figures 1b-d, individuals in red are those within the *H* subsample and individuals in blue are those excluded from the *H* subsample (i.e., the subsample with the smallest determinant). Figure 1b shows the observed squared MD (horizontal) and *χ*^2^-distances (vertical). The *χ*^2^-distances are the squared component scores (*cf*. Eq. (10)). We have denoted two individuals with “X” circumscribed by a circle. These two individuals are the only two with “aa” for one particular SNP. Finally, we show the robust squared MD, which helps reveal “masking” effects: Figure 1c shows the observed vs. robust squared MD. Note the substantial increase of the two previously denoted outliers. Figure 1d is the same as Figure 1d except without the two most extreme individuals to provide a better view of the observed vs. robust squared MDs.

**Figure 1.**
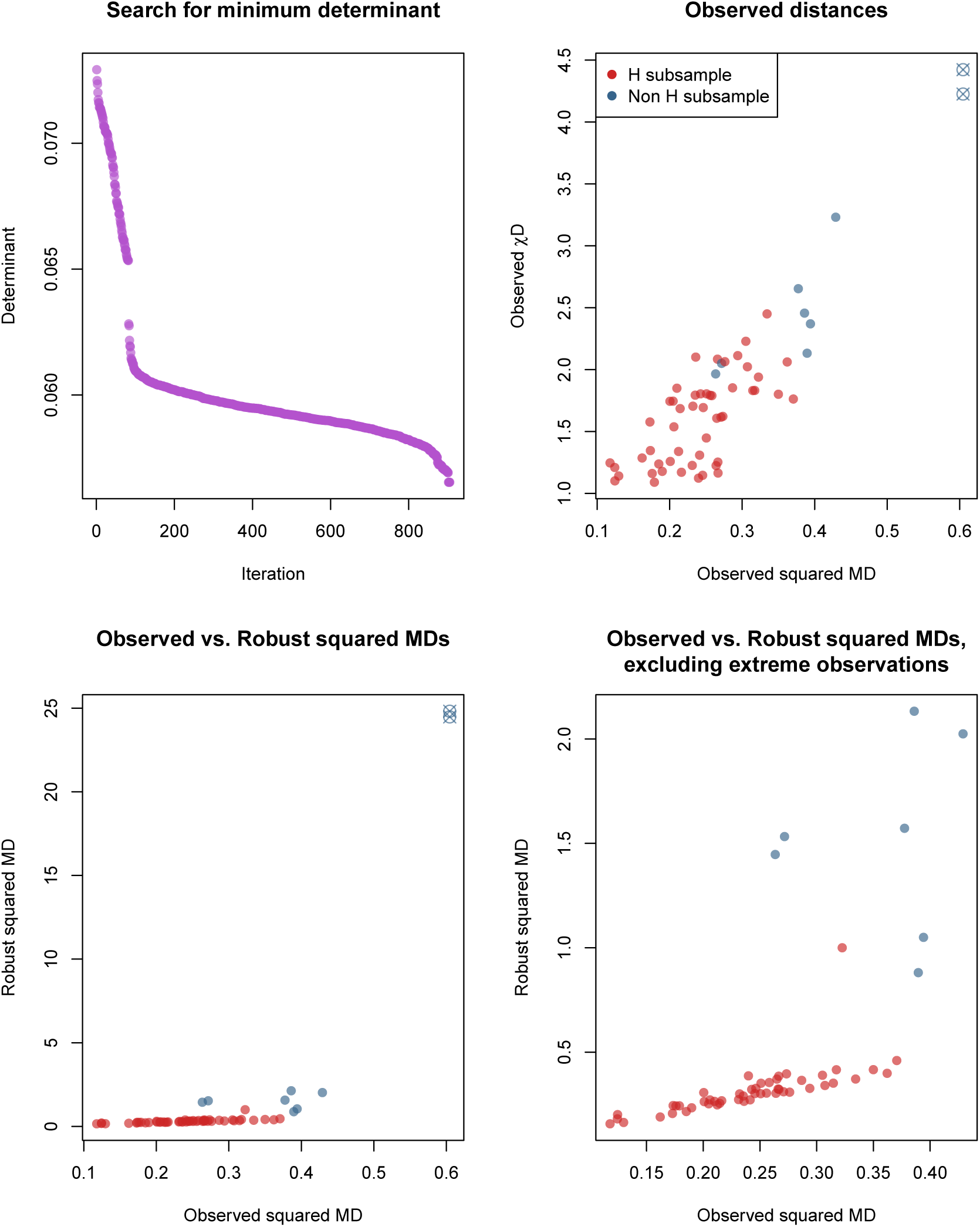
Application of generalized MCD to categorical data. We illustrate the generalized MCD here first on a small data set of single nucleotide polymorphisms (SNPs). (a) shows the search for the determinant. (b) shows the observed distances with the final *H* subset colored in red. (c) shows the observed vs. robust squared Mahalanobis distances (MDs). (d) shows the same as (c) but excludes the two extreme individuals (which were the only two to have ‘aa’ for one particular SNP.

### ONDRI Data

The Ontario Neurodegenerative Disease Research Initiative (ONDRI; http://ondri.ca/) is a longitudinal, multi-site, “deep-phenotyping” study across multiple neurodegenerative disease cohorts (Farhan et al., 2017). The data presented here are preliminary data, and used for illustrative purposes. As part of the quality control process, various subsets of data are subjected to an outlier analyses pipeline (Sunderland et al., 2019) via the MCD and CorrMax (Garthwaite & Koch, 2016) procedures. In ONDRI, many individual variables or entire instruments are not quantitative, such as the MSAM. The MSAM is a short version of the survey of autobiographical memory (SAM). The MSAM is comprised of 10 questions where each question has five possible responses: two responses indicate disagreement with a question (“strongly disagree”, “disagree somewhat”), two responses indicate agreement with a question (“agree somewhat”, “strongly agree”) and one neutral respone (“neither”). Here we apply two strategies to the data—as categorical, then as ordinal—on *N* = 300 from two cohorts (cerebrovascular disease *n* = 161 and Parkinson’s disease *n* = 139). We compare and contrast the results of the two strategies and at three *α* levels: *α* = 0.5, *α* = 0.75, *α* = 0.9 (for a total of six applications of the GMCD).

First, we treat the data as categorical and apply our technique as previously outlined. Next, we use a specific recoding scheme for ordinal data which has many names: “bipolar coding” (Greenacre, 1984), fuzzy coding (Goldfarb & Pardoux, 2006), or “doubling” (Greenacre, 2014; Lebart et al., 1984). We refer to the doubling approach for ordinal data as “fuzzy coding” (described below). Fuzzy coding transforms variables into “pseudo-disjunctive” in that, from the perspective of CA and MCA, the data tables behave like disjuntive tables (see Table 1): the sums of the rows equal the number of (original) variables, the sum of variables (each pair of columns) equal the number of rows, and the sum of the table equals the number of rows × the number of (original) variables. With fuzzy coding, we represent each original variable with two columns that reflect the distance between an ordinal response and its expected minimum or expected maximum. The pair of columns for each variable, referred to here as “−” and “+”, sum to 1. So 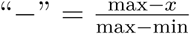 reflects how close to the minimum of the scale the observed (*x*) value is, and 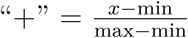 reflects how close to the maximum of the scale the observed (*x*) value is. The MSAM has 5 possible levels (1, 2, 3, 4, 5), thus our expected—but not necessarily observed—minimum and maximum for each response is 1 and 5, respectively. We illustrate fuzzy coding in Table 2.

**Table 2.** Example of fuzzy coding.

|  | Q1: Specific events | Q2: Recall objects |
| --- | --- | --- |
| *OND01_TWH_1100* | 2 | 4 |
| *OND01_SBH_1200* | 1 | 4 |
| ... | ... | ... |
| *OND01_HDH_5100* | 5 | 4 |
| *OND01_HDH_5201* | 3 | 5 |

(b) *Doubling for ordinal data ("fuzzy coding") with normalization so each variable sums to 1*
|  | Q1− | Q1+ | Q2− | Q2+ |
| --- | --- | --- | --- | --- |
| *OND01_TWH_1100* | $\frac{5-2}{5-1} = 0.75$ | $\frac{2-1}{5-1} = 0.25$ | $\frac{5-4}{5-1} = 0.25$ | $\frac{4-1}{5-1} = 0.75$ |
| *OND01_SBH_1200* | $\frac{5-1}{5-1} = 1.00$ | $\frac{1-1}{5-1} = 0.00$ | $\frac{5-4}{5-1} = 0.25$ | $\frac{4-1}{5-1} = 0.75$ |
| ... | ... | ... | ... | ... |
| *OND01_HDH_5100* | $\frac{5-5}{5-1} = 0.00$ | $\frac{5-1}{5-1} = 1.00$ | $\frac{5-4}{5-1} = 0.25$ | $\frac{4-1}{5-1} = 0.75$ |
| *OND01_HDH_5201* | $\frac{5-3}{5-1} = 0.50$ | $\frac{3-1}{5-1} = 0.50$ | $\frac{5-5}{5-1} = 0.00$ | $\frac{5-1}{5-1} = 1.00$ |

*Note.* Example of data doubling via fuzzy coding for ordinal data with the miniature survey of autobiographical memory (MSAM). Note that the sum of each row is equal to the total number of original variables, and that the sum within each pair of columns per variable is equal to the total number of rows.

As in the Fast-MCD algorithm, *α* controls the proportion of samples to identify as the *H* subset. The size of *H* is computed as *H* = ⌊(2 × *H*_*I*_) − *I* + (2 × (*I* − *H*_*I*_) × *α*) ⌋ where *H*_*I*_ = mod ((*I* + *J* + 1), 2) (which is the same subsample size computation as the `rrcov` and robustbase packages in `R` via the `h.alpha.n` function). Because the categorical and the ordinal versions of the data sets have a different number of columns (*J*) after transformation into disjunctive or pseudo-disjunctive data, the same *α* produced a different size *H* for each data set. Each application of these analyses had an upper limit of 1, 000 iterations to search for a minimum determinant.

Figure 2 shows all six applications of the generalized MCD applied to the MSAM. To note in Figure 2 we show the MDs and robust MDs (not the squared MDs or robust MDs), because this better highlights the differences on a smaller scale for the axes. Generally, when *α* is low, we see greater exaggeration of outlying individuals with respect to the “inliers” and the *H* subset. However this behavior essentially disappears as *α* increases. We also see that the “ordinal” version of the data behaves quite differently from the categorical version, especially as *α* increases. In particular, we see considerable changes in the robust MDs for the individuals excluded from the *H* subset.

**Figure 2.**
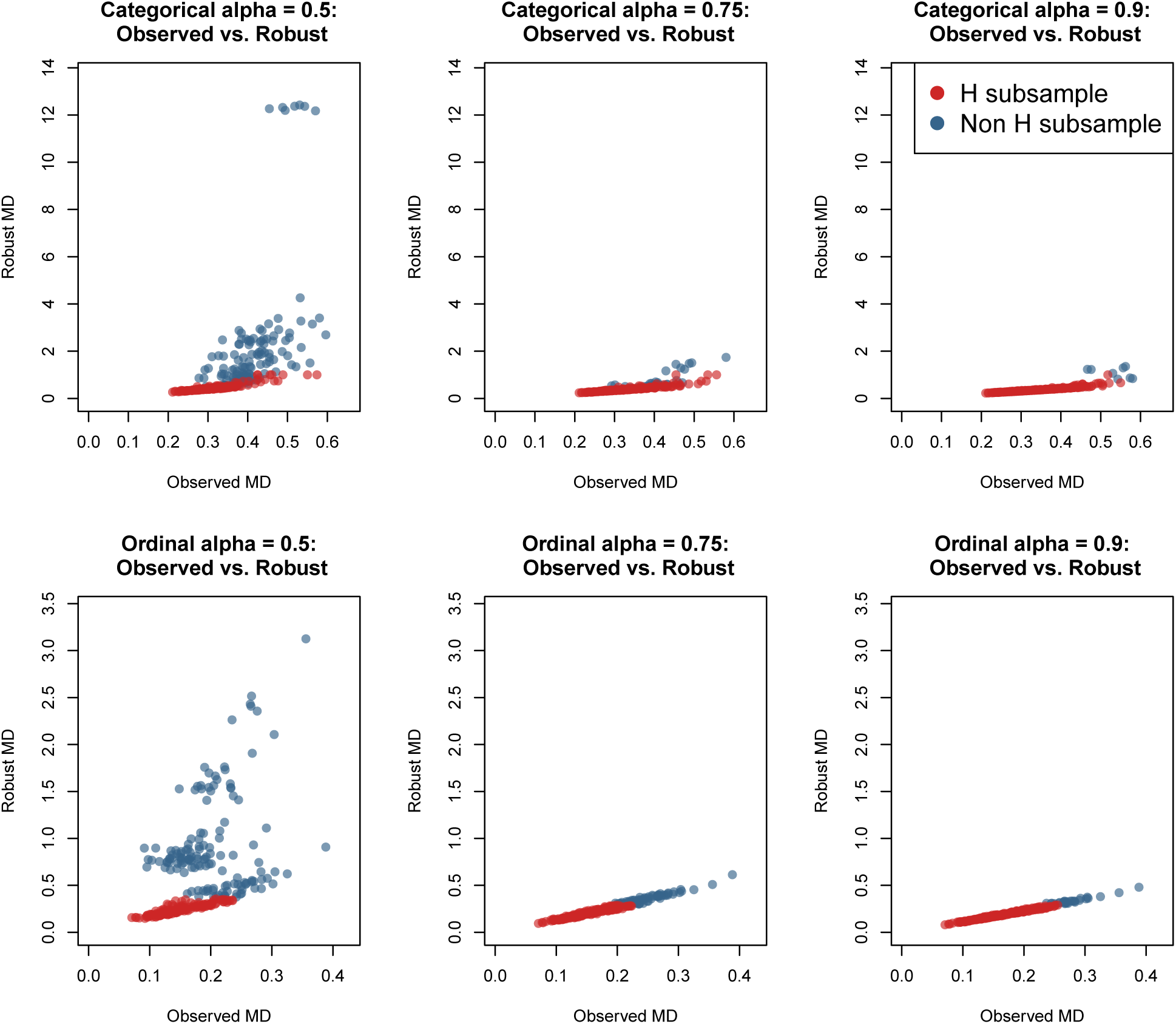
Generalized MCD applied to the miniature survey of autobiographical memory (MSAM) data from ONDRI. Left to right is the alpha level and top to bottom is how the data were coded. All plots show observed vs. robust Mahalanobis distances (MD). Observations in red are the H subsample, observations in blue are not in the *H* subsample. Axes are scaled to the same values within each recoding to highlight the changes in robust MD.

There are differences in how the robust MDs express themselves with respect to the robust covariance structure in each analysis. We can see that the categorical version exaggerates the robust MD across all versions, which highlights the masking effect. We still see this effect in the ordinal version, but only with the lower *α* level – once *α* is increased, the robust and standard MDs are quite similar. However, even though the robust MDs express varying patterns (see Fig. 2), the identification of the *H* subsamples across the different analyses are nearly identical. The overlap of the *H* subsamples (see Table 3) shows that each approach to the data (categorical or ordinal) identifies many of the same *H* subsample. The main difference between the categorical and ordinal approaches is the magnitude of the robust MDs with respect to their robust covariance structures.

**Table 3.** Generalized MCD subsample sizes.

|  | C .5 | C .75 | C .9 | O .5 | O .75 | O .9 |
| --- | --- | --- | --- | --- | --- | --- |
| C .5 | 175 |  |  |  |  |  |
| C .75 | 165 | 237 |  |  |  |  |
| C .9 | 168 | 228 | 275 |  |  |  |
| O .5 | 127 | 149 | 155 | 160 |  |  |
| O .75 | 159 | 205 | 223 | 150 | 230 |  |
| O .9 | 172 | 229 | 262 | 160 | 230 | 272 |

*Note.* C = Categorical, O = Ordinal. Only the lower triangle and diagonal are shown. The diagonal indicates the size of the H subset out of $N = 300$ . Off-diagonal values indicate the overlap in subsets between each version of the miniature survey on autobiographical memory (MSAM) analysis. As the proportion for subsample sizes increase, overlap becomes more likely.

### ADNI Data

Data used in the preparation of this article come from Phase 1 of the ADNI database (adni.loni.usc.edu). ADNI was launched in 2003 as a public-private funding partnership and includes public funding by the National Institute on Aging, the National Institute of Biomedical Imaging and Bioengineering, and the Food and Drug Administration. The primary goal of ADNI has been to test a wide variety of measures to assess the progression of mild cognitive impairment and early Alzheimer’s disease. The ADNI project is the result of efforts of many coinvestigators from a broad range of academic institutions and private corporations. Michael W. Weiner (VA Medical Center, and University of California-San Francisco) is the ADNI Principal Investigator. Subjects have been recruited from over 50 sites across the United States and Canada (for up-to-date information, see www.adni-info.org).

We used three data sets from ADNI: SNPs (categorical data), the clinical dementia rating (CDR; as ordinal data), and volumetric estimates from five brain regions (continuous data). The ADNI sample for use here was *N* = 613 across three diagnostic groups: Alzheimer’s disease (AD) = 264, Mild cognitive impairment (MCI) = 166, and a control group (CON) = 183. These data can be obtained in ADNI via the genome-wide data and the `ADNIMERGE R` package.

We used SNPs associated with APOE and TOMM40, which are strong genetic contributions to AD (Roses et al., 2010). SNPs were excluded if their minor allele frequency was < 5%. Participants or SNPs with *>* 5% missingness were excluded. Genotypes with < 5% were recoded; generally the minor homozygote (“aa”) had low frequency therefore in those cases we coded for the presence or absence of “a” (i.e., “AA” vs. presence of “a”; which is the dominant inheritance model). We were thus left with 13 SNPs that spanned 35 columns: 4 of our SNPs were recoded to combine “Aa” and “aa” (i.e., “AA” vs. “Aa+aa”) and the remaining 9 SNPs each had three levels (i.e., “AA”, “Aa”, and “aa”).

The CDR is a structured interview that has six domains: memory, orientation, judgement, communication, home and hobbies, and self-care (Morris (1993); see also http://alzheimer.wustl.edu/cdr/cdr.htm). After the interview ratings are applied to each category with the following possible responses: 0 (normal), 0.5 (very mild), 1 (mild), 2 (moderate), and 3 (severe). The CDR has ordinal responses and therefore we recoded the CDR with fuzzy coding.

We also included five brain volumetric estimates known to be impacted by AD: ventricles, hippocampus, enthorinal cortex, fusiform gyrus, and medial temporal regions. The volumetric estimates here are treated as continuous data. We used a “data doubling” approach for continuous data so that it too behaves like disjunctive data. We refer to the transformation of continuous to pseudo-disjunctive data specifically as the “Escofier transform” (Beaton et al., 2016; Escofier, 1979). The Escofier transform has a lower and upper value like fuzzy coding: 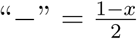 and 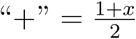, where *x* is typically a normalized form of a continuous value (e.g., a *Z*-score). An example of “doubling” for continuous data can be seen in Table 4.

**Table 4.**
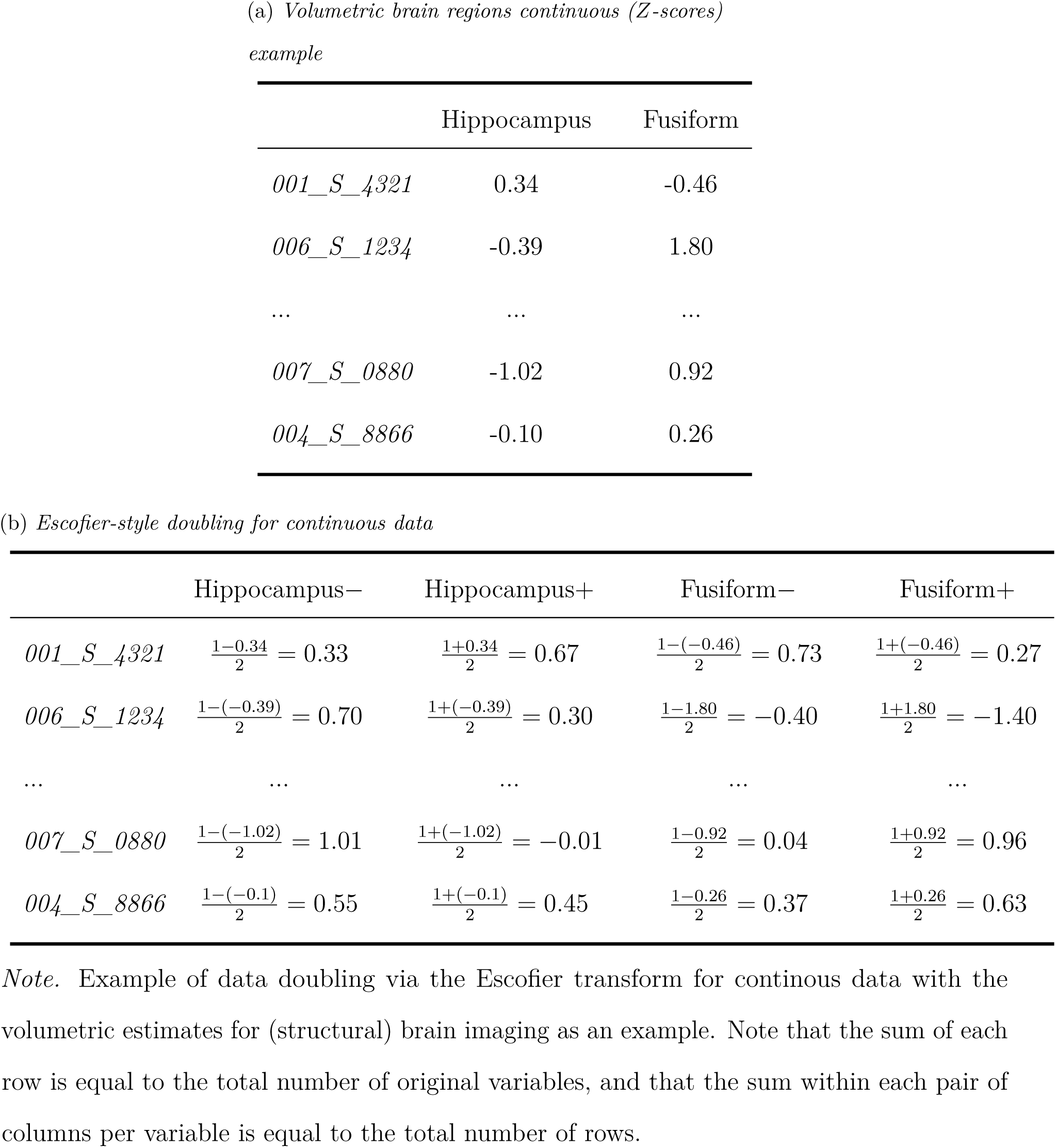
Example of Escofier transform.

*Volumetric brain regions continuous (Z-scores)example*
|  | Hippocampus | Fusiform |
| --- | --- | --- |
| *001_S_4321* | 0.34 | -0.46 |
| *006_S_1234* | -0.39 | 1.80 |
| ... | ... | ... |
| *007_S_0880* | -1.02 | 0.92 |
| *004_S_8866* | -0.10 | 0.26 |

(b) *Escofier-style doubling for continuous data*
|  | Hippocampus− | Hippocampus+ | Fusiform− | Fusiform+ |
| --- | --- | --- | --- | --- |
| *001_S_4321* | $\frac{1-0.34}{2} = 0.33$ | $\frac{1+0.34}{2} = 0.67$ | $\frac{1-(-0.46)}{2} = 0.73$ | $\frac{1+(-0.46)}{2} = 0.27$ |
| *006_S_1234* | $\frac{1-(-0.39)}{2} = 0.70$ | $\frac{1+(-0.39)}{2} = 0.30$ | $\frac{1-1.80}{2} = -0.40$ | $\frac{1+1.80}{2} = -1.40$ |
| ... | ... | ... | ... | ... |
| *007_S_0880* | $\frac{1-(-1.02)}{2} = 1.01$ | $\frac{1+(-1.02)}{2} = -0.01$ | $\frac{1-0.92}{2} = 0.04$ | $\frac{1+0.92}{2} = 0.96$ |
| *004_S_8866* | $\frac{1-(-0.1)}{2} = 0.55$ | $\frac{1+(-0.1)}{2} = 0.45$ | $\frac{1-0.26}{2} = 0.37$ | $\frac{1+0.26}{2} = 0.63$ |

*Note.* Example of data doubling via the Escofier transform for continuous data with the volumetric estimates for (structural) brain imaging as an example. Note that the sum of each row is equal to the total number of original variables, and that the sum within each pair of columns per variable is equal to the total number of rows.

After preprocessing we had a matrix with *I* = 613 ×*J* = 57 (13 SNPs categorical that spanned 35 columns, 6 CDR domains that spanned 12 columns, and 5 brain regions that spanned 10 columns). In this example, we set *α* = 0.5. Figure 3 shows our GMCD applied to the mixed data set, where individuals in red are those included in the *H* subsample and individuals in blue are those excluded from the *H* subsample. Figure 3a shows the observed squared MD and *χ*^2^-distances for individuals (in red) that comprise the minimum determinant. Figure 3b shows the observed vs. robust MD. Figure 3c shows the log transformed observed vs. robust MD to help visualize the entire set of individuals. In Figure 3c we also include a horizontal line to help denote which individuals were susceptible to the “masking” effect.

**Figure 3.**
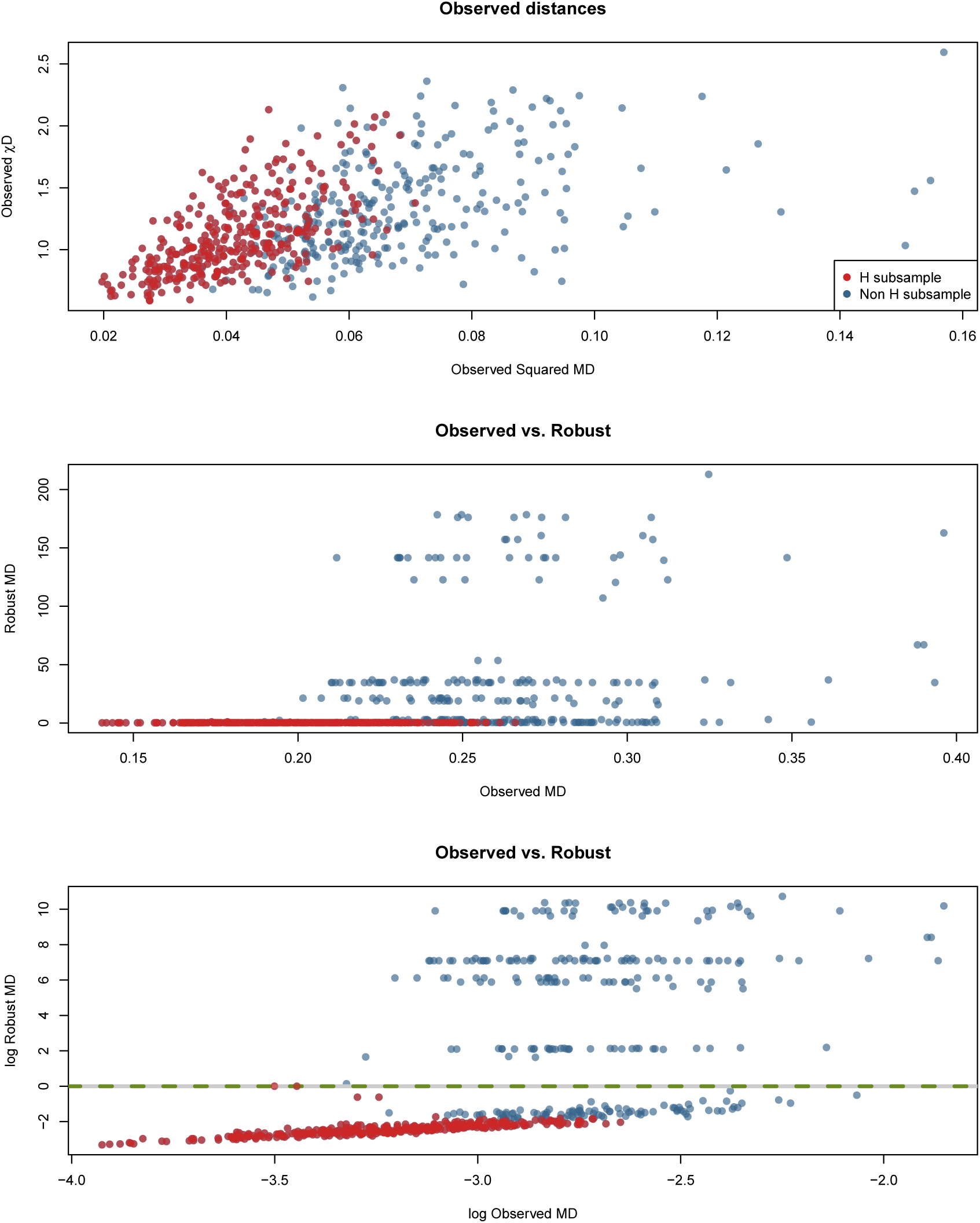
Application of generalized MCD to mixed data. Data are comprised of SNPs (categorical), clinical dementia rating (ordinal) and volumetric brain estimates (continuous) from ADNI. (a) shows the observed distances with the *H* subset colored in red, where (b) shows the observed vs. robust Mahalanobis distances (MDs), and (c) shows the same but with log transformed observed and robust distances. Panel (c) includes a horizontal line to highlight individuals susceptible to the ‘masking’ effect.

While MCD algorithms are useful to identify outlying individuals in a multivariate space, these algorithms do not indicate why individuals are outliers. To help understand why these individuals are outliers we used a heatmap of the 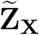 matrix (Figure 4). The rows in Figure 4 are the observations where the columns are the variables. For the columns, we excluded the “lower pole” from the data-doubling procedure of the ordinal and continuous values as these are redundant in the visualization.

**Figure 4.**
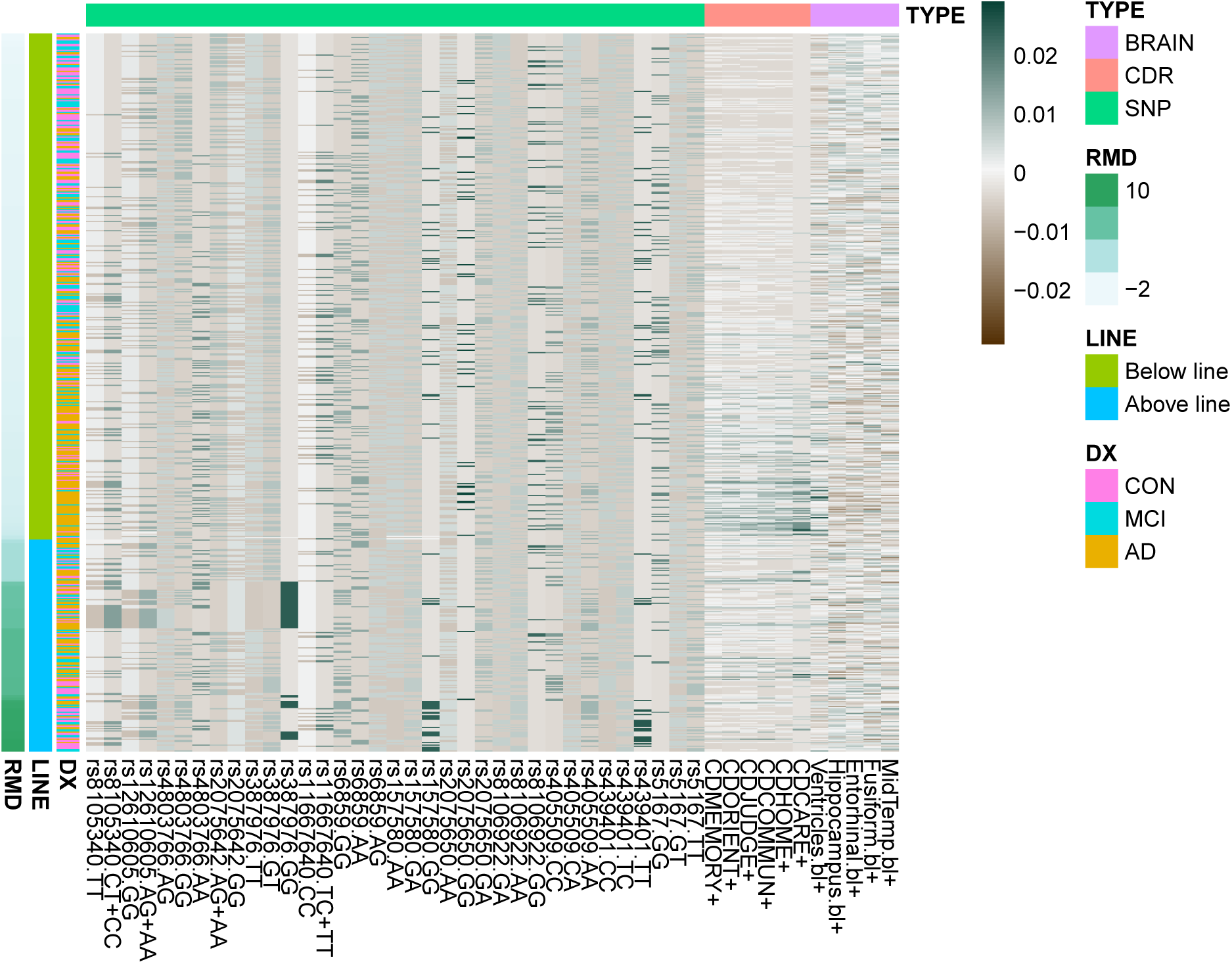
A heatmap of the 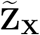 for the ADNI data. The scale is based on the maximum absolute value in 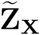. TYPE denotes which type of data each were in each column. To note, here we only show the ‘upper pole’ of the clinical dementia rating (CDR; ordinal) and brain volume (continuous) data. Rows of the heatmap are in descending order of robust Mahalanobis distance. The rows have three labels: (1) RMD which is the log transform of the robust Mahalanobis distance (as seen in the previous figure), (2) LINE which denotes which rows are above or below the horizontal line in the previous figure, and (3) DX which is the diagnosis of each participant.

The scale in Figure 4 comes from the maximum absolute value from the 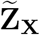 matrix. For the columns, we indicate which data type each column belongs to with color in the legend—brain volumes (continuous), clinical dementia rating (ordinal), and SNPs (categorical). For the rows, we provide three separate labels: (1) a continuous label that is the (natural) log transform of the robust MDs, (2) whether the row (observation) is above or below the horizontal line in Figure 3c, and (3) the diagnostic label of each participant where CON = control, MCI = mild cognitive impairment, and AD = Alzheimer’s disease. We generally refer to the individuals “above” the line as outliers in this case.

The outliers in Figure 4 do not show a homogeneous pattern. That is, there exist multiple ways in which an individual could be an outlier. Generally, when compared to the “inliers” (those below the line) there exists clear patterns of outlierness. First, a particular genotype—the GG genotype in SNP rs387976—exists exclusively within the outlier set. However, not all individuals within the outlier set have the GG genotype in SNP rs387976. In the outlier set, we can see that there also exists patterns of rare genotypes. For example, quite a few individuals have both the the GG genotype for rs157580 and TT for rs439401. But there also exists additional patterns in the outlier set that are unexpected. Generally, as clinical symptoms of AD present, we also expect that ventricle volumes increase, while the other brain volumes decrease. Instead, we see individuals that have those brain patterns with low CDR ratings (i.e., no clinical indiciation of AD). We also see the opposite: higher CDR ratings with brain volume patterns that do not reflect atrophy or degeneration. In some cases, we see only specific high (or low) values for the CDR and brain volumes, which indicate specific unexpected values as opposed to unexpected patterns. Furthermore, some individuals in the outlier set express unique patterns across all measures, e.g., individuals with infrequent genotypes and unexpected high (or low) CDR scores with low (or high) brain volumes. The diagnostic label provides critical information. The outlier set is not comprised of any single diagnosis. Rather, the outlier set is diagnostically heterogeneous, which means that these individuals are atypical with respect to the entire sample and especially for their respective diagnostic groups.

## Discussion

We presented a MCD-based approach that generalizes to virtually any data type (i.e., categorical, ordinal, continuous). The basis of our approach relies on Mahalnobis distances from CA via the SVD. Our definition of MDs avoids the issues—such as excessive weights and arbitrary metrics—of Mahalanobis-like distances that Goodall (1966) noted. And because our generalized MCD relies on CA, this allows for various types or mixtures of data through simple recoding approaches (see Tables 2 and 4). Our generalized MCD was designed around the necessity to identify robust structures and outliers in complex non-quantitative data. And like many MCD-based and -inspired approaches (Boudt et al., 2017; Fritsch, Varoquaux, Thyreau, Poline, & Thirion, 2012; Mejia et al., 2017; Rousseeuw & Van Driessen, 1999), our approach identifies both robust substructures and outliers (often hidden through “masking” effects).

Within the ONDRI study, we realized that many—if not all—existing outlier detection and robust covariance techniques could not accomodate our needs. We required a multivariate outlier technique that could easily work on categorical, ordinal, or mixed data. While there are other possible algorithms (e.g., Candès, Li, Ma, & Wright, 2011; Fan, Liao, & Mincheva, 2013) we chose to extend the MCD because it is a reliable and widely-used technique (Rousseeuw & Van Driessen, 1999 has been cited 2, 238 times at the time we wrote this paper), and available in languages routinely used for multivariate analyses (e.g., https://cran.r-project.org/web/packages/robustbase/index.html, https://cran.r-project.org/web/packages/rrcov/index.html, and https://wis.kuleuven.be/stat/robust/LIBRA). We have made our GMCD approach available in an R package designed to detect outliers and robust structures (see https://github.com/derekbeaton/ours).

### Future directions and applications of GMCD

In our ADNI example we treated each variable as an independent measure. Though routine, that is somewhat unrealistic: each individual variable comes from larger sets of related variables (e.g., SNPs, the CDR, and brain imaging). Multiple Factor Analysis (MFA; Escofier & Pages, 1994; Abdi, Williams, & Valentin, 2013) is a technique designed to accomodate *a priori* groupings of variables (columns) with respect to a global analysis. Notably, MFA has been extended to mixtures of data types (Bécue-Bertaut & Pagès, 2008) based on the work by Escofier (1979) and Escofier and Pages (1994) that we use here. Thus, our GMCD approach could be applied for MFA-type problems.

Because our generalized MCD is based on GCA, as defined by Escofier (1983) and Escofier (1984), we do not depend on the 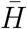 sample set to 0s in the iterative process. The use of GCA allows for any reasonable definition of weights applied to the rows or columns, as well as defining alternate expected values. For example, we could use population level estimates of the allele frequencies (from, e.g., dbGaP, HapMap or 1000 Genomes), which would change our column weights and expected matrix. Likewise, we could also use alternate row weights. In standard MCA—and here in our GMCD—all observations are assigned equal weights (i.e., 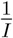). But in standard CA, rows do not necessarily have equal weights. Rather the row weights are proportional to all elements in that row akin to the column weights (*cf*. Eq. (6)). Though these variations are possible, we have not explored them here and consider these as open questions.

As previously noted, in MCA we discard null dimensions (i.e., those with effectively 0 eigenvalues). However, in MCA it is also typical to discard many of the low-variance components with eigenvalues (Abdi & Valentin, 2007). In MCA, the reason to discard these low-variance components is because of a correction to the dimensionality of the space, which should reflect the *C* original variables as opposed to the *J* analyzed columns. The “Benzécri correction” is the most common approach to identify which low-variance components to discard (Benzécri, 1979), which are components with eigenvalues smaller than 1*/C*. The Benzécri correction suggests a way to retain a lower dimensional space, and also means we can compute Mahalanobis-like distances from some arbitrary set of high-variance components. This would be equivalent to a rudimentary regularization. However in our GMCD approach, we do not discard components via Benzécri correction, rather we retain all non-null dimensions for analyses.

### Limitations

While our approach opens many avenues to extend the MCD there are some limitations. As just noted, the Benzécri correction (Benzécri, 1979) discards low-variance components. First, the Benzécri correction does not work well in practice for use in the GMCD. In most cases outliers typically express high values on low-variance components. If we were to discard low-variance components we might not identify a suitable robust structure nor identify outliers. The low variance components are typically where the masking effect exists. Furthermore, the dimensionality correction (Benzécri, 1979) of the space is only established for strictly categorical—and thus completely disjunctive—data. How to determine, and then use, the correct dimensionality of the space (especially for mixed variables) is an open question. We have opted for the more conservative approach: require full rank data *I > J* and retain all components with non-zero eigenvalues.

As noted in our concluding points in our formulation of the GMCD, the use of multiple columns to represent each variable means there exists linear dependency between the columns within any given variable. By definition, the determinant of such a matrix is 0. Our GMCD solution relaxes the requirement of a determinant and makes use of all non-null dimensions from MCA, and thus is a pseudo-determinant. Likewise, our MD is also defined only from those non-null dimensions.

Finally, our algorithm does not compute a robust center from the subsample. To do so would likely drop levels from variables (effectively zeroing out columns). When discarding specific columns, the MDs of individuals with those levels *decreases*. This decrease is incorrect, because often those levels are rare, and thus the individuals should have a high MD. Rather, our robust center exists within the *χ*^2^-space as defined by the factor scores. This is because subset MCA “maintains the geometry […] and *χ*^2^-distances of the complete MCA […]” (Greenacre & Pardo, 2006). For example if we were to apply our technique *as is* with continuous data (recoded as in Table 4) and compare against the standard MCD we would obtain slightly different results. However, at each step if we were to re-center the continuous data before applying the Escofier transform, we would produce exactly the same results as the standard MCD.

Note that we also do not provide any discussion on thresholds to identify outliers. In most applications of the MCD, a cutoff values is defined as some percentage (e.g., 97.5%) and relies on various distributions such as *χ*^2^ (Hubert et al., 2017) or the *F*-distribution (Hardin & Rocke, 2005). Rather, we encourage the use of resampling methods and to use empirical cutoffs from the data set. This could be done with distributions of robust distances generated from, for examples, bootstrap or repeated subsampling techniques.

### Conclusions

The MCD is a reliable and widely-used approach to identify robust multivariate structures and outliers. Until now, the MCD could only be used for data that are (or were assumed to be) continuous. Our approach uses CA to define Mahalanobis distances via the singular vectors and allows us to apply an MCD approach to virtually any data type. We believe that our generalized MCD via GCA opens many new avenues to develop multivariate robust and outlier detection approaches for the complex data sets we face today. Not only does GCA provide the basis for a new family of MCD techniques but our technique also suggests ways to approach other robust techniques (Candès et al., 2011; Fan et al., 2013). With studies such as ONDRI and ADNI, the boundaries of what constitutes a “data set” are fluid and now inherently heterogeneous: researchers analyze multiple variables of different types from various sources. Our examples highlight these types of routine data sets. With complex studies that contain many mixed-variables, techniques like our generalized MCD will be essential to maintain data quality and integrity.

## ONDRI

This work was completed on behalf of the Ontario Neurodegenerative Disease Research Initiative (ONDRI). The authors would like to acknowledge the ONDRI Founding Authors: Robert Bartha, Sandra E. Black, Michael Borrie, Dale Corbett, Elizabeth Finger, Morris Freedman, Barry Greenberg, David A. Grimes, Robert A. Hegele, Chris Hudson, Anthony E. Lang, Mario Masellis, William E. McIlroy, Paula M. McLaughlin, Manuel Montero-Odasso, David G. Munoz, Douglas P. Munoz, J.B. Orange, Michael J. Strong, Stephen C. Strother, Richard H. Swartz, Sean Symons, Maria Carmela Tartaglia, Angela Troyer, and Lorne Zinman.

## ADNI

Data collection and sharing for this project was funded by the Alzheimer’s Disease Neuroimaging Initiative (ADNI) (National Institutes of Health Grant U01 AG024904) and DOD ADNI (Department of Defense award number W81XWH-12-2-0012). ADNI is funded by the National Institute on Aging, the National Institute of Biomedical Imaging and Bioengineering, and through generous contributions from the following: AbbVie, Alzheimer’s Association; Alzheimer’s Drug Discovery Foundation; Araclon Biotech; BioClinica, Inc.; Biogen; Bristol-Myers Squibb Company; CereSpir, Inc.; Cogstate; Eisai Inc.; Elan Pharmaceuticals, Inc.; Eli Lilly and Company; EuroImmun; F. Hoffmann-La Roche Ltd and its affiliated company Genentech, Inc.; Fujirebio; GE Healthcare; IXICO Ltd.; Janssen Alzheimer Immunotherapy Research & Development, LLC.; Johnson & Johnson Pharmaceutical Research & Development LLC.; Lumosity; Lundbeck; Merck & Co., Inc.; Meso Scale Diagnostics, LLC.; NeuroRx Research; Neurotrack Technologies; Novartis Pharmaceuticals Corporation; Pfizer Inc.; Piramal Imaging; Servier; Takeda Pharmaceutical Company; and Transition Therapeutics. The Canadian Institutes of Health Research is providing funds to support ADNI clinical sites in Canada. Private sector contributions are facilitated by the Foundation for the National Institutes of Health (www.fnih.org). The grantee organization is the Northern California Institute for Research and Education, and the study is coordinated by the Alzheimer’s Therapeutic Research Institute at the University of Southern California. ADNI data are disseminated by the Laboratory for Neuro Imaging at the University of Southern California.

